# Compositionally distinct nuclear pore complexes of functionally distinct dimorphic nuclei in ciliate *Tetrahymena*

**DOI:** 10.1101/116277

**Authors:** Masaaki Iwamoto, Hiroko Osakada, Chie Mori, Yasuhiro Fukuda, Koji Nagao, Chikashi Obuse, Yasushi Hiraoka, Tokuko Haraguchi

## Abstract

**SUMMARY STATEMENT:** Our study demonstrates compositional and structural differences of the nuclear pore complex between the functionally differentiated macronucleus and micronucleus within a single cytoplasm of ciliated protozoa.

**ABSTRACT:** The nuclear pore complex (NPC), a gateway for nucleocytoplasmic trafficking, is composed of about 30 different proteins called nucleoporins. It remains unknown whether the NPCs within a species are homogeneous or vary depending on the cell type, or physiological condition. Here, we present evidence for compositionally distinct NPCs that form within a single cell in a binucleated ciliate. In *Tetrahymena thermophila,* each cell contains both a transcriptionally-active macronucleus (MAC) and a germline micronucleus (MIC). By combining *in silico* analysis, mass spectrometry analysis for immuno-isolated proteins, and subcellular localization analysis of GFP fused proteins, we identified numerous novel components of MAC and MIC NPCs. Core members of the Nup107-160 scaffold complex were enriched in MIC NPCs. Strikingly, two paralogs of Nup214 and of Nup153 localized exclusively to either MAC or MIC NPCs. Furthermore, the transmembrane components Pom121 and Pom82 localize exclusively to MAC and MIC NPCs, respectively. Our results argue that functional nuclear dimorphism in ciliates is likely to depend on compositional and structural specificity of NPCs.

## INTRODUCTION

Ciliated protozoa maintain two distinct nuclei within the same cytoplasm: a somatic macronucleus (MAC) and a germline micronucleus (MIC) (Fig. 1A) (Eisen et al., 2006; Orias et al., 2011; Karrer, 2012). The polyploid MAC is transcriptionally active, and its acentromeric chromosomes segregate during cell division by a spindle-independent amitotic process. In contrast, the diploid MIC has transcriptionally inert, centromeric chromosomes that segregate by canonical mitosis. In *Tetrahymena thermophila,* DNA replication in the MIC and MAC occurs during non-overlapping periods in the cell cycle. Thus, nuclear dimorphism in ciliates involves non-equivalent regulation of multiple activities in two distinct nuclei (Orias, 2000; Goldfarb and Gorovsky, 2009). This is likely to require targeted transport of components to the MIC *vs.* MAC, for which differences in the NPCs may be important determinants.

Previously, we analyzed 13 *Tetrahymena* nucleoporins (Nups), and discovered that four paralogs of Nup98 were differentially localized to the MAC and MIC (Iwamoto et al., 2009). The MAC- and MIC-specific Nup98s are characterized by Gly-Leu-Phe-Gly (GLFG) and Asn-Ile-Phe-Ans (NIFN) repeats, respectively, and this difference is important for the nucleus-specific import of linker histones (Iwamoto et al., 2009). The full extent of compositional differentiation of MAC and MIC NPCs could not, however, be assessed, since only a small subset of the expected NPC components were detected.

NPCs have been studied in eukaryotes including rat (Cronshaw et al., 2002), *Saccharomyces cerevisiae* (Rout et al., 2000), *Aspergillus nidulans* (Osmani et al., 2006), *Schizosaccharomyces pombe* (Asakawa et al., 2014), *Arabidopsis thaliana* (Tamura et al., 2010), and *Trypanosoma brucei* (Degrasse et al., 2009; Obado et al., 2016) (Table S1). The NPC structure has an 8-fold rotational symmetry, and is made up of roughly 30 known Nups organized into subcomplexes (Alber et al., 2007; Bui et al., 2013) (Fig. S1). The Nup93 complex in mammalian cells (Nic96 in *S. cerevisiae*) forms a stable scaffold composed of Nup93^*Sc*Nic96^, Nup205^*Sc*Nup192^, Nup188^*Sc*Nup188^, Nup155^*Sc*Nup170 or *Sc*Nup157^, and Nup53/Nup35/MP-44^*Sc*Nup53 or *Sc*Nup59^ (Grandi et al., 1997; Hawryluk-Gara et al., 2005; Amlacher et al., 2011). A second stable scaffold in mammals, the Nup107-160 complex (called the Y-complex or Nup84 complex in *S. cerevisiae*) is composed of conserved subunits Nup107^*Sc*Nup84^, Nup160^*Sc*Nup120^, Nup133^*Sc*Nup133^, Nup96^*Sc*Nup145C^, Nup85^*Sc*Nup85^, Seh1, and Sec13, together with species-specific subunits (Siniossoglou et al., 1996; Lutzmann et al., 2002; Loiodice et al., 2004). Peripheral to the scaffolds are Phe-Gly (FG) repeat-bearing Nups, whose disordered FG-repeat regions constitute the central channel, with FG repeats interacting with nuclear transport receptors (Terry and Wente, 2009). Three transmembrane (TM) Nups anchoring the NPC to the mammalian nuclear membrane are NDC1, gp210, and POM121 (Greber et al., 1990; Hallberg et al., 1993; Stavru et al., 2006) (in yeast: Ndc1, Pom152, and Pom34 (Winey et al., 1993; Wozniak et al., 1994; Miao et al., 2006)). A distinct nucleoplasmic basket is formed with Tpr^*Sc*Mlp1/Mlp2^ (Cordes et al., 1997; Strambio-de-Castillia et al., 1999).

Based on prior analysis, *T. thermophila* appeared to lack homologs of many widely conserved NPC components. These included scaffold Nups (mammalian Nups205, 188, 160, 133, 107, 85, and 53, among others) from the Nup93 and Y-complexes. Similarly, homologs of FG-Nups Nup214, 153, 62, and 58 were also not detected, as were TM Nups except for gp210. These NPC components may have evaded homology-based searches due to extensive sequence divergence, given the large evolutionary distance between ciliates and animals, fungi, and plants.

To address these ambiguities and to better understand NPC differentiation in *T. thermophila,* we attempted comprehensive identification of Nups. First, we analyzed proteins affinity-captured with known Nups. Furthermore, we mined updated genome and protein databases for characteristic Nup sequences or conserved domains, using *in silico* structure prediction. The resulting expanded catalog of *Tetrahymena* Nups, combined with localization data, sheds new light on the extent to which NPC architecture can vary within a single species, and even in a single cytoplasm.

## RESULTS

### The Nup93 complex includes a unique Nup205 ortholog and a novel central channel FG-Nup

In mammalian cells, the Nup93 complex (Fig. 1B) is composed of Nup93, Nup205, Nup188, Nup155, and Nup53 (Fig. S1) (Grandi et al., 1997; Hawryluk-Gara et al., 2005). In *Tetrahymena,* we previously identified homologs for Nup93 (T*t*Nup93; Gene Model identifier TTHERM_00622800) and Nup155 (T*t*Nup155; TTHERM_00760460), and found them distributed to MAC and MIC NPCs (Iwamoto et al., 2009). To identify other Nup93 complex components, we used mass spectrometry to analyze anti-GFP immunoprecipitates from *Tetrahymena* expressing GFP-T*t*Nup93 (Fig. 1C). All of the proteins listed in Table S2 as ‘hypothetical protein’ were examined by Blast search for similarities to known Nups of other species. In addition, all of the ‘hypothetical proteins’ were examined by expression profile analysis in the *Tetrahymena* Functional Genomics Database (TetraFGD) web site (http://tfgd.ihb.ac.cn/) (for details see the “Microarray” page of the TetraFGD: http://tfgd.ihb.ac.cn/tool/exp (Miao et al., 2009)) (also see Materials and Methods). When either the Blast search or the expression profile analysis (details described below) found similarities to any known Nups, we examined its subcellular localization in *T. thermophila* by ectopically expressing GFP fused proteins. By these analyses we found Nup308 (TTHERM_00091620) and the novel protein TTHERM_00194800 (T*t*Nup58: Nup58 in Fig. 1D and Table S2).

**Fig. 1.**
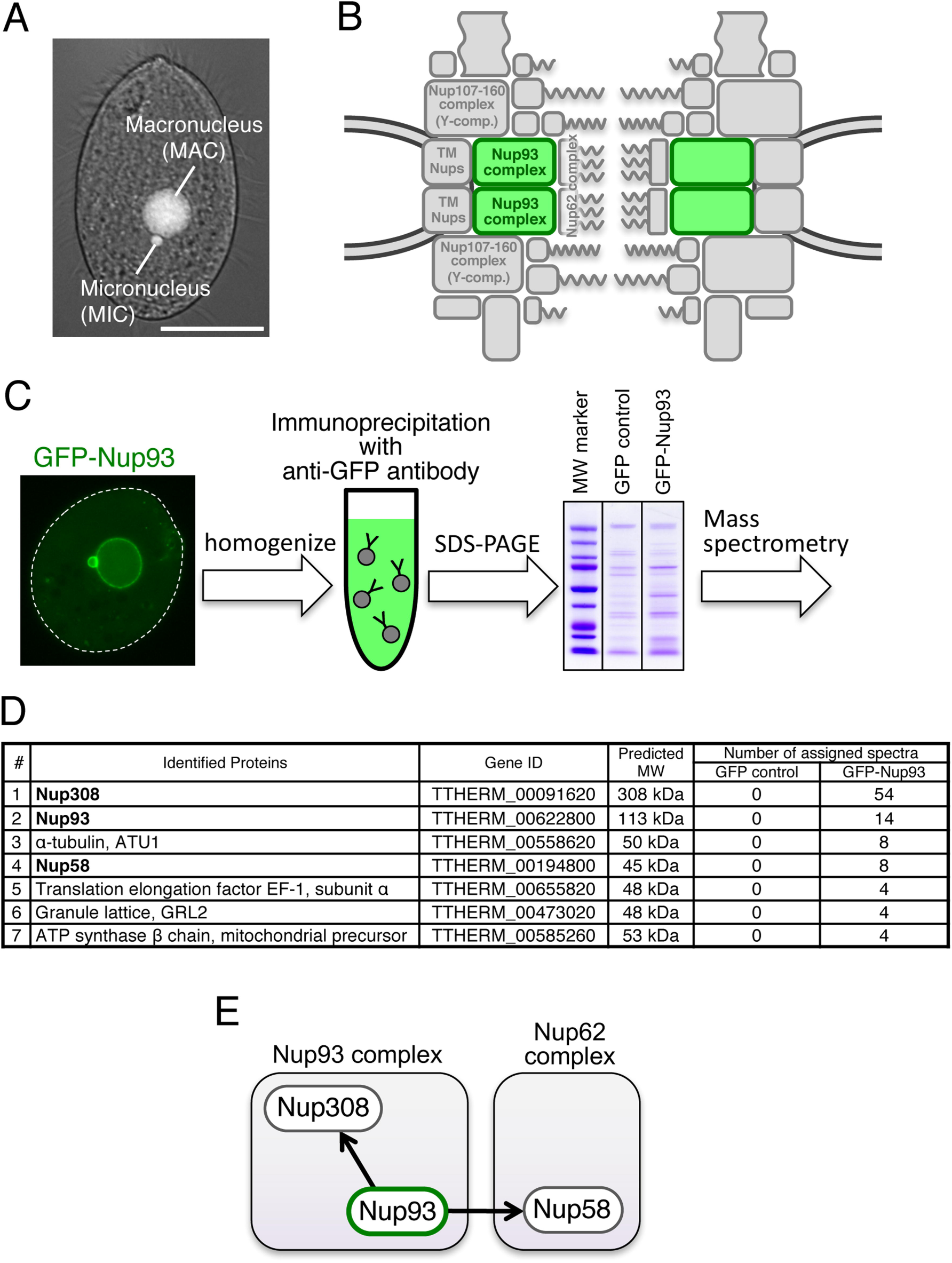
Immunoprecipitation and mass spectrometry analysis to identify Nup93 complex members. (A) A *T. thermophila* cell fixed with methanol and stained with DAPI. Bar, 20 μm. (B) The position of the Nup93 complex within the NPC architecture. See also Fig. S1. (C) Simplified procedure of immunoprecipitation and mass spectrometry for GFP**–***Tt*Nup93-expressing cells used for immunoprecipitation. (D) Mass spectrometric identification of the proteins co-precipitated with GFP**-***Tt*Nup93. The top seven proteins are listed among other identified proteins (Table S2). (E) Physical interaction map of Nup93 based on the mass spectrometry results.

Nup308, a protein of 2675 amino acid residues, was previously identified as a *Tetrahymena-*specific Nup, but it was not assigned to a subcomplex (Iwamoto et al., 2009). Based on PSIPRED analysis, Nup308 is composed of GLFG repeats forming an N-terminal disordered structure (residues 1–570), followed by a large C-terminal α-helix-rich region (residues 571–2675) (Fig. 2). To identify potential Nup308 counterparts, we looked for Nups in other species with similar distributions of secondary structures. Interestingly, a large α-solenoid domain is a predicted feature of both Nup205 and Nup188, conserved core members of the Nup93 complex (Kosova et al., 1999; Andersen et al., 2013), although these proteins do not have FG repeats.

**Fig. 2.**
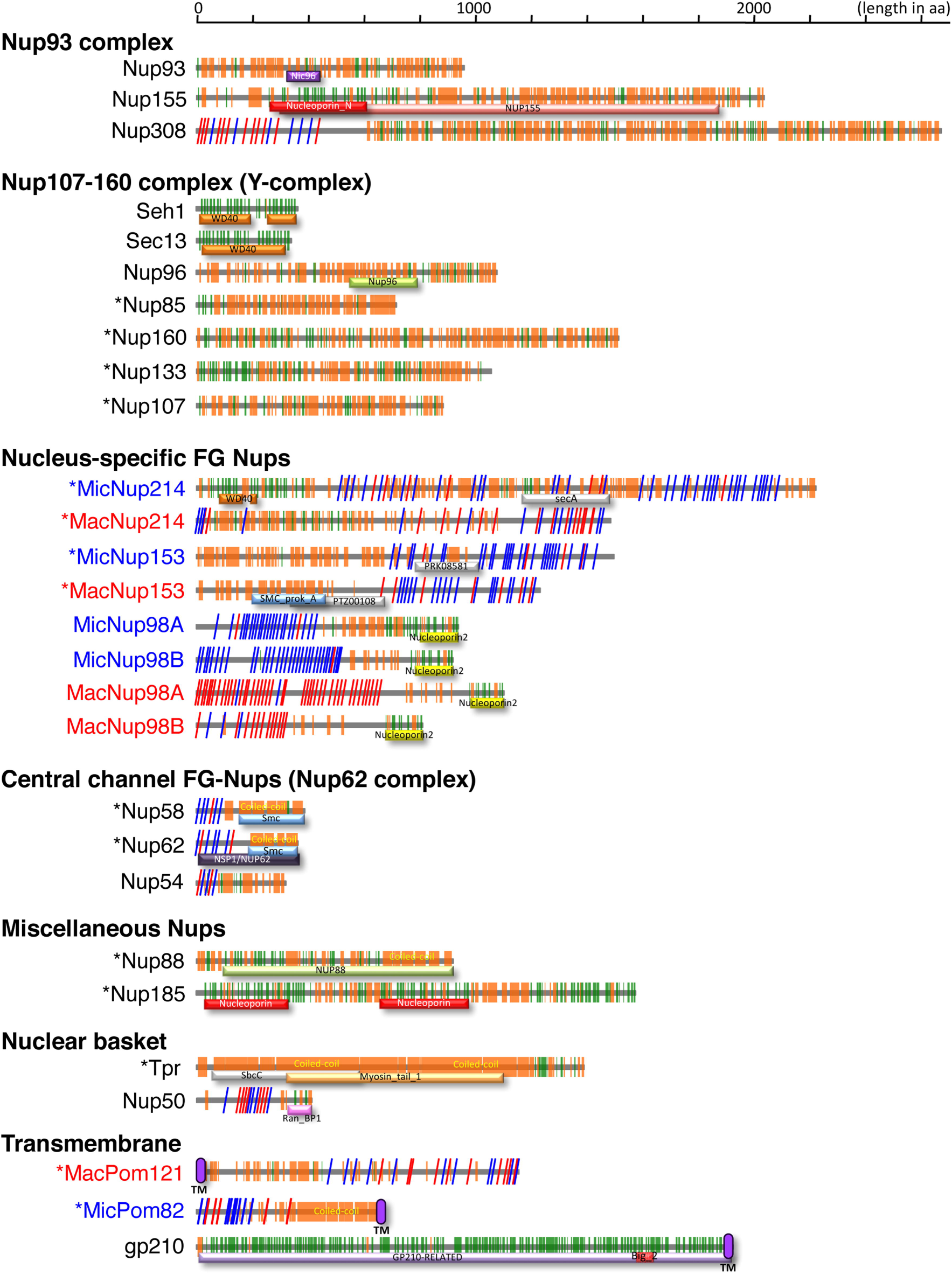
Distributions of secondary structures and conserved domains in *Tetrahymena* nucleoporins. Each Nup is shown as the protein name on the left. Blue, red, and black letters mean MIC-specific, MAC-specific, and shared components, respectively. Asterisks on the left shoulder of the protein names indicate Nups newly identified in this study. The colored components in the illustration are as follows: orange boxes/bars, α-helix; green boxes/bars, β-strand; red slanting lines, FG repeats; blue slanting lines, FX repeats (X means any residue, but the majority are N and Q); and purple ellipse, predicted TM domain. Conserved domains are indicated by differently colored bars with standard domain names.

To investigate whether this structural similarity between *Tetrahymena* Nup308 and Nup205 and Nup188 homologs in other species reflected shared evolutionary origins, we performed a phylogenetic analysis. Nup308 formed a clade with Nup205 orthologs, supported by a bootstrap probability of 72%, but not with Nup188 orthologs (Fig. S2). Nup188 appears absent in *Tetrahymena,* since we failed to find any candidates in either the database or in our mass spectrometry data. Taken together, our results strongly suggest that Nup308 belongs to the Nup93 complex and is orthologous to human Nup205, but has acquired an unusual GLFG repeat domain. Consistent with this assignment, GFP-Nup308 localized similarly to GFP-*Tt*Nup93, being equally distributed between MAC and MIC NPCs (Iwamoto et al., 2009).

The second Nup candidate identified in *Tt*Nup93 pulldowns was TTHERM_00194800. This small protein (45 kDa deduced molecular weight) is composed of an N-terminal FG-repeat region and a C-terminal coiled-coil region (Fig. 2), which are characteristics of central channel FG-Nups that are tethered by Nup93 (Chug et al., 2015). The secondary structure characteristics of the novel *Tetrahymena* Nup are highly similar to those of Nup62 and Nup58, central channel proteins in yeast and vertebrates that interact with Nup93 (Grandi et al., 1993, 1997). Because another protein was found as an Nup62 ortholog (described below), this protein is the likely *Tetrahymena* ortholog of Nup58; therefore, we named it *Tt*Nup58 (Nup58 in Fig. 1D,E).

### Newly identified members of the Y-complex are likely homologs of conserved Nups

The vertebrate’s Y-complex (Fig. 3A) contains 10 distinct proteins (Orjalo et al., 2006; Mishra et al., 2010), of which 3 had identified homologs in *Tetrahymena* (*Tt*Seh1, *Tt*Sec13, *Tt*Nup96) (Iwamoto et al, 2009). To investigate whether the remaining seven are present in *Tetrahymena* but had been overlooked due to sequence divergence, we carried out mass spectrometric analysis of anti-GFP immunoprecipitates from cells expressing the known Y-complex GFP-tagged Nups described below.

First, in precipitates of GFP–*Tt*Seh1, we identified an 86 kDa protein orthologous to Nup85 (Table S3) with a short stretch of four predicted β-strand blades at the N-terminus followed by an α-solenoid domain (Fig. 2). That architecture is typical of Nup85 orthologs that are Y-complex components in other organisms (Brohawn et al., 2008). We therefore tentatively named the *Tetrahymena* protein *Tt*Nup85. GFP–TtNup85 localized to NPCs in both the MAC and MIC (Figs 3B and S3A).

**Fig. 3.**
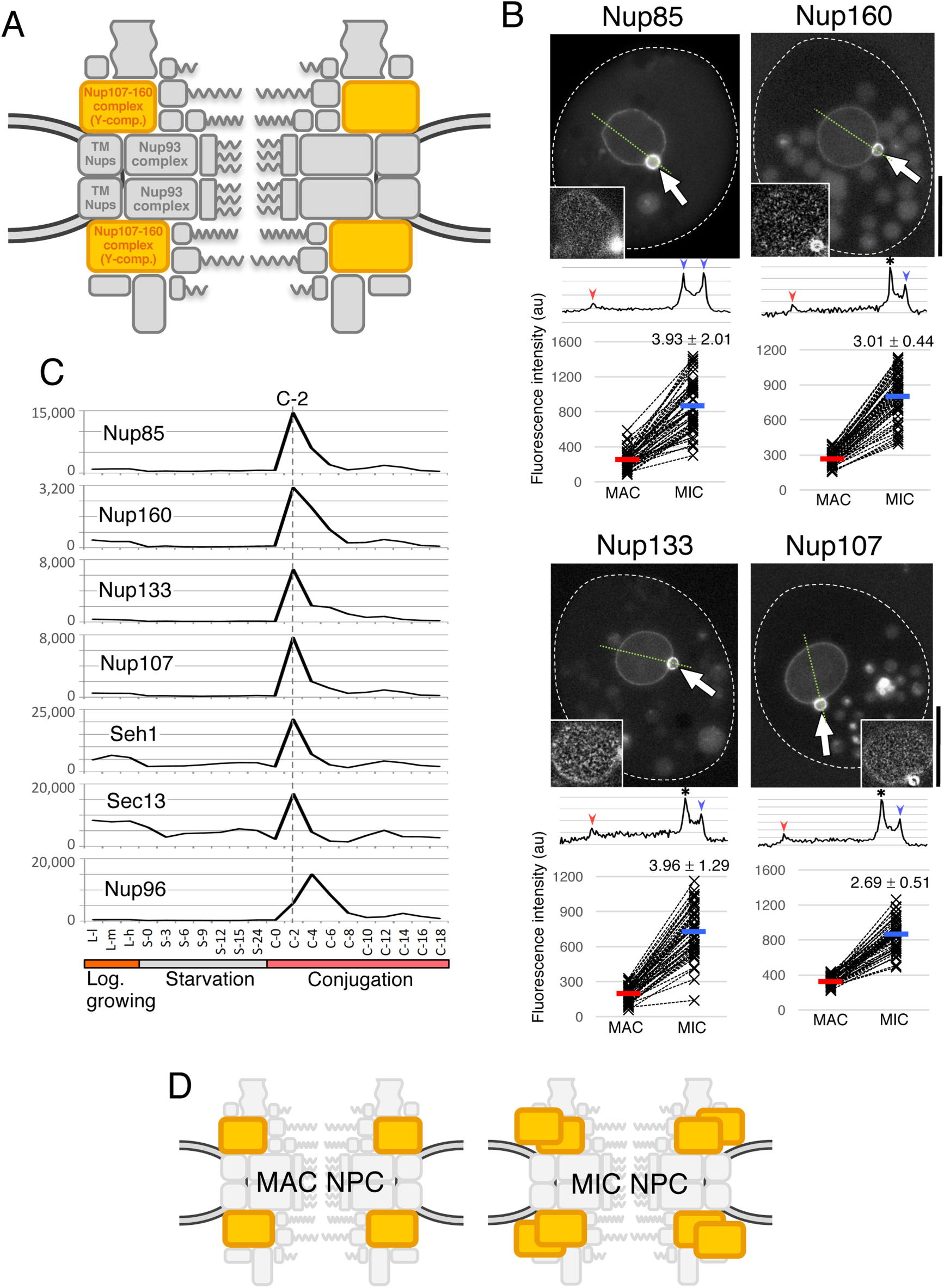
Y-complex components localize to both nuclei but are biased to MICs. (A) The position of the Y-complex within the NPC architecture. (B) Fluorescent micrographs of GFP-Nups ectopically expressed in *Tetrahymena* cells. White broken lines represent the borders of cells. The inset in each panel shows a deconvoluted image focused on the MAC surface. Arrows indicate the position of the MIC. Bars, 20 μm. A line profile of fluorescence intensity along the thin green broken line is presented under each image panel. Blue and red arrowheads indicate the points corresponding to MIC and MAC envelopes, respectively. An asterisk marks the point at which the borders of the two nuclei overlap, and where the intensity is measured as the sum of both NEs. Below the line profile, the fluorescence intensities of MAC and MIC NEs from 50 cells are plotted. The vertical axis of the graph is shown in arbitrary units. Broken lines connect the plots of MAC and MIC within the same cell. Average values are presented by red and blue bars for MAC and MIC, respectively. The numbers upon the MIC plots indicate fold increase of fluorescence in MIC from MAC. All differences are significant (*P* < 10^−20^ by Student’s *t*-test). (C) Expression profiles of the Y-complex members extracted from the TetraFGD (http://tfgd.ihb.ac.cn/). Plots are the average of two values presented in the database. The horizontal axis represents successive stages of culture growth and therefore different physiological conditions. For the logarithmic growth stage, L-l, L-m, and L-h represent low, medium, and high cell concentrations, respectively. For starvation and conjugation stages, numbers represent hours after the transfer of the cells to each condition. The vertical axis represents relative values of mRNA expression. For details, visit the database website. (D) A simple representation of the deduced composition of MAC and MIC NPCs with different numbers of Y-complexes.

We then immunoprecipitated GFP–*Tt*Nup85, and identified two novel candidate Y-complex core members. Both proteins are composed of a β-strand-rich N-terminal half and an α-helical-rich C-terminal half. This domain architecture is characteristic of the Y-complex components Nup160 and Nup133 (Berke et al., 2004; Devos et al., 2004), and we tentatively named the *Tetrahymena* proteins *Tt*Nup160 and *Tt*Nup133 (Fig. 2 and Table S4). GFP–*Tt*Nup160 and GFP–*Tt*Nup133 localized to NPCs in both nuclei, like other Y-complex components (Figs 3B and S3A).

Another conserved Y-complex component is Nup107, which interacts with Nup96. To search for the *Tetrahymena* homolog we used GFP–*Tt*Nup96 as bait and identified a 109 kDa protein (Table S5) that is rich in predicted α-helices like human Nup107 (Fig. 2). The protein, tentatively named *Tt*Nup107, localized as a GFP-tagged construct to NPCs of both nuclei (Figs 3B and S3A).

The genes encoding all members of the Y-complex except for Nup96 are co-expressed and exhibit sharp expression peaks at 2 h (C-2) after two cell strains with different mating-types were mixed for conjugation (for details see the “Microarray” page of the TetraFGD: http://tfgd.ihb.ac.cn/tool/exp (Miao et al., 2009)) (Fig. 3C). In contrast, *Tt*Nup96 exhibits an expression peak at 4 h (C-4). This difference in the timing of expression between *Tt*Nup96 and the other Y-complex components may be related to a unique aspect of *Tt*Nup96 gene structure: *Tt*Nup96 is expressed as part of a single transcription unit together with MicNup98B, under the promoter of the MicNup98B gene (Iwamoto et al., 2009).

Three other components of the human Y-complex were not detected in our studies: Nup43, Nup37, and ELYS. These components may be species-specific (Bilokapic and Schwartz, 2012; Rothballer and Kutay, 2012), and genuinely absent from *Tetrahymena.* They are also absent from *S. cerevisiae* (Alber et al., 2007) (see Table S1), supporting this idea.

### Y-complex components show biased localization to the MIC

As previously reported, GFP-tagged Nup93 complex members and some of the central channel Nups (*Tt*Nup93, *Tt*Nup308, and *Tt*Nup54) were distributed equally between MAC and MIC NPCs, judging by fluorescence intensities (Iwamoto et al., 2009). In striking contrast, all Y-complex components so far identified exhibit distinctively biased localization to the MIC nuclear envelope (NE) compared to the MAC NE (Fig. 3B). Fluorescence intensities in the MIC were 2.69–3.96 times higher than those in the MAC (Fig. 3B). This biased localization of Y-complex components may be caused by overexpression of the components due to ectopically expressing the GFP-tagged proteins in addition to the expression of endogenously untagged ones. To address this issue, we examined the localization of Nup160-GFP, Nup133-GFP, and Seh1-mCherry expressed from endogenous loci under the control of their native promoters, and therefore expressed at physiological levels. All three proteins showed biased localization, as found for the overexpressed GFP-tagged proteins (compare the images in Fig. 3B and Fig. S3B), suggesting that the biased localization is not caused by overexpression of the tagged proteins. Because the NPC density is similar in the MAC and MIC (Fig. S1 in Iwamoto et al. (2009)), the relative concentration of Y-complex components in the MIC NE suggests that the Y-complex is present at higher copy number per NPC in the MIC compared to the MAC (Fig. 3D).

### Newly-detected FG-Nups include nucleus-specific and common components

FG-Nups were originally characterized as nucleoporins with domains containing extensive repeats of phenylalanine-glycine (FG) that function in nucleocytoplasmic transport. More recently, we reported a remarkable difference in MAC and MIC NPCs regarding the repeat signature present in four Nup98 paralogs. The repeat signature of MacNup98A and -B is mostly GLFG, while that of MicNup98A and -B is mostly NIFN (Fig. 2) (Iwamoto et al., 2009; 2010; 2015). We have now taken advantage of the recently improved annotation of the *Tetrahymena* Genome Database Wiki (http://ciliate.org/index.php/home/welcome), to search for sequences bearing repeats that are similar to those of FG-Nups in other species. We found five candidate FG-Nups. Based on the molecular size and the positions of predicted α-helices, β-strands, and FG-repeat regions, we designated four of these proteins as MicNup214 (TTHERM_00992810), MacNup214 (TTHERM_00755929), MicNup153 (TTHERM_00647510), and MacNup153 (TTHERM_00379010): GFP-fusions of MicNup214 and MacNup214 were exclusively localized to the MIC and MAC, respectively (Fig. 4A,B). Fluorescent protein (GFP or mNeon)-fusions of MicNup153 were mostly localized to the MIC and secondarily to the MAC in most growing cells (Fig. 4A), although it was exclusively localized to the MIC in some cells (Fig. S3C). GFP-fusions of MacNup153 were exclusively localized to the MAC (Fig. 4B). The localization of the fifth candidate FG-Nup (*Tt*Nup62: Nup62 in Fig. 4C), like the novel nucleoporin *Tt*Nup58 (Nup58 in Fig. 4C) identified as a central channel protein (discussed above), was less specific.

A striking feature of the Nup214 paralogs is that they contain the same nucleus-specific repeat motifs described earlier for *Tt*Nup98 paralogs. Like the MIC-specific Nup98 paralogs, MicNup214 contains NIFN repeats (the last N is mostly Q in this protein), while MacNup214 contains FG repeats (Fig. 2). This difference may be an important determinant for selective protein transport to the MAC and MIC, as previously shown for *Tt*Nup98s (Iwamoto et al., 2009). We note that MacNup214 lacks a β-strand-rich N-terminal region that is found in other Nup214 orthologs (Weirich et al., 2004; Napetschnig et al., 2007) (Fig. 2).

In contrast, MicNup153 and MacNup153 do not differ markedly from one another in their molecular features (Fig. 2). Because the N-terminus domain of human Nup153 is involved in its NPC localization (Enarson et al., 1998), we speculate that the N-terminal domains of MicNup153 and MacNup153 may also be involved in their nucleus-specific localization in *Tetrahymena*. Further study is required to elucidate their nucleus-specific localization.

While the expression of this set of FG-Nups is upregulated during conjugation (Fig. 4D), the MIC-specific components tend to be expressed 2 h earlier than MAC-specific ones. For example, MicNup214 expression peaks at 2 h in conjugation (C-2) vs. MacNup214 at C-4; similarly, MicNup153 peaks at C-6 vs. MacNup153 at C-8 (Fig. 4D). The earlier expression of MIC-specific components compared with MAC-specific ones may reflect a selective requirement for MIC-specific NPCs during early stages of conjugation, such as the crescent stage (Sugai and Hiwatashi, 1974). In contrast, the later expression of MAC-specific components probably reflects formation of the new MACs that occurs in the later stages of conjugation.

**Fig. 4.**
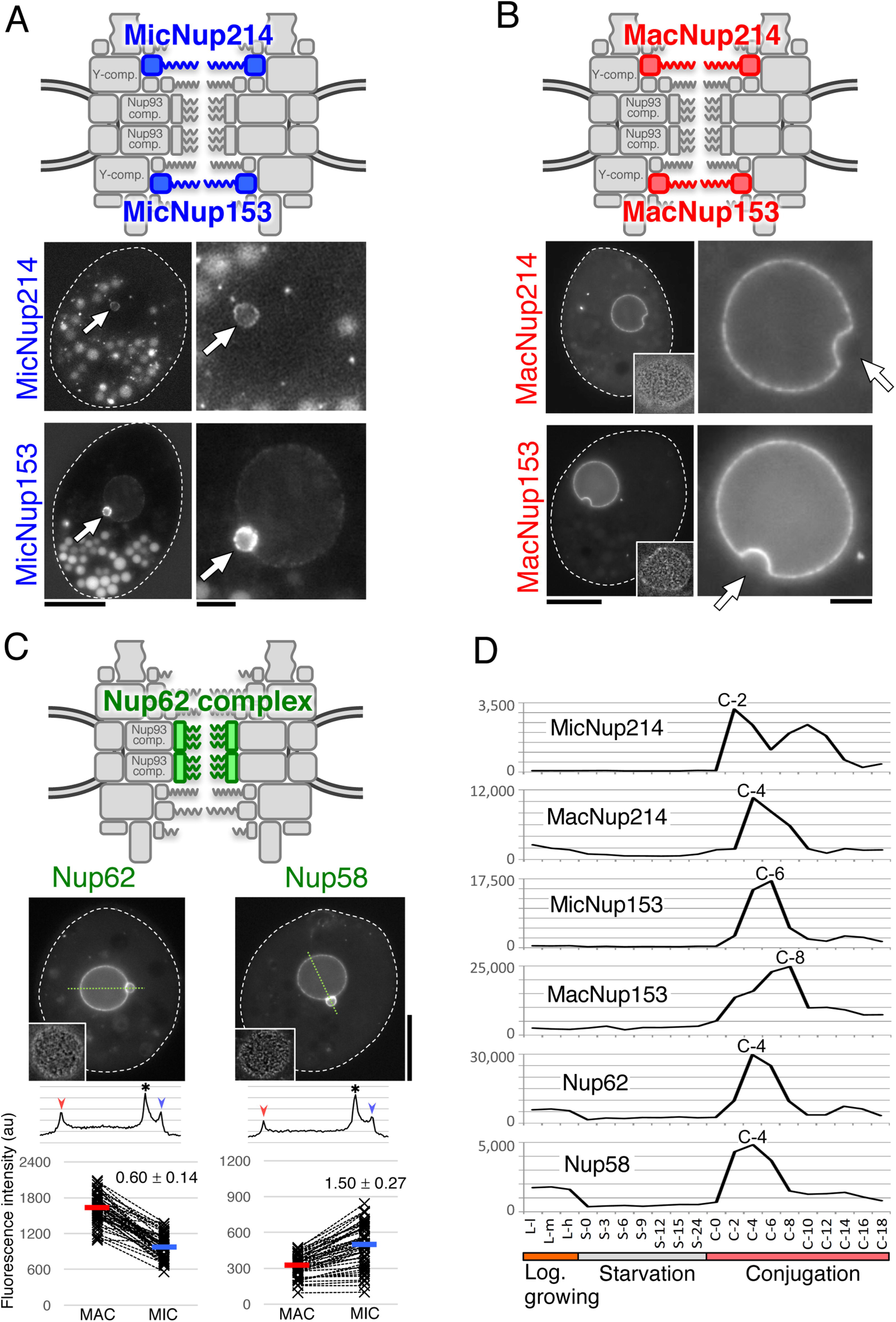
Newly identified FG-Nups of *Tetrahymena.* (A) MIC-specific paralogs of Nup214 and Nup153. The upper figure indicates the predicted positions of these Nups within the MIC NPC. Fluorescence micrographs show the subcellular localization of fluorescent protein-tagged Nups; MicNup214 and MicNup153 were endogenously tagged with GFP and mNeon at the C-termini of their ORFs, respectively. Arrows indicate the position of the MIC. Other fluorescent bodies dispersed in the cytoplasm are phagosomes taking in materials derived from the culture medium. (B) MAC-specific paralogs of Nup214 and Nup153. The upper figure indicates the predicted positions of these Nups within the MAC NPC. Fluorescence micrographs show the subcellular localizations of ectopically expressed GFP**–**Nups. The left panels show a whole cell, and each nuclear region is enlarged in the right panels. White broken lines represent the borders of cells. Insets in the left panels show deconvoluted images focused on the MAC surface. Arrows indicate the position of MICs. Bars indicate 20 μm for the left panels and 5 μm for the right panels. (C) *Tt*Nup62 and *Tt*Nup58 localized in both nuclei. The upper illustration indicates the predicted position of these Nups, which constitute the Nup62 complex. Fluorescent micrographs show the subcellular localizations of ectopically expressed GFP**–***Tt*Nup62 and *Tt*Nup58**–**GFP. Bars, 20 μm. Line profiles and plots of fluorescence intensity are shown under each image panel in the same manner as in Fig. 3B. Both differences are significant (*P* < 10^−16^ by Student’s *t*-test). (D) Expression profiles of FG-Nups, as in Fig. 3C.

The fifth candidate FG-Nup identified by this screen was a 39 kDa protein (TTHERM_01122680). This protein is composed of an N-terminal FG-repeat region and a C-terminal coiled-coil region with the characteristics of central channel FG-Nups and is assigned as a Nucleoporin NSP1/NUP62 family protein (IPR026010) (Fig. 2). Consequently, this protein is the likely *Tetrahymena* ortholog of Nup62; therefore, we named it *Tt*Nup62. The GFP-tagged protein was distributed to both nuclei (Nup62 in Fig. 4C), similarly to the central channel Nups *Tt*Nup58 (Figs 1E and 4C) and *Tt*Nup54 (Iwamoto et al., 2009), although *Tt*Nup62 was slightly enriched in the MAC NE, whereas *Tt*Nup58 was slightly enriched in the MIC NE. The expression profile of *Tt*Nup62 was similar to that of *Tt*Nup58, with an expression peak after 4 h of conjugation (C-4) (Fig. 4D).

*Tt*Nup62 has relatively few repeats in its FG motif compared with homologs such as human Nup62 and *S. cerevisiae* Nsp1 (Fig. 2), although it has several FX repeats (X=N, Q, A or T in the case of this protein). A feature unique to *Tetrahymena* is the presence of GLFG repeats in Nup308, an ortholog of Nup205. The Nup93 complex containing Nup205 anchors Nup62 (Vollmer and Antonin, 2014), and it is likely that the *Tetrahymena* Nup93 complex containing Nup308 anchors *Tt*Nup62. Thus, we hypothesize that the GLFG repeats present in Nup308 compensate for the low number of FG repeats of *Tt*Nup62 presents in the central channel.

### Nup88, Nup185, and Tpr

We used a variety of strategies to identify additional Nups. Homology searches against InterPro (http://www.ebi.ac.uk/interpro/) revealed a gene (TTHERM_00455610) with a conserved Nup88 domain “*Tt*Nup88 (PTHR13257:SF0)” (Fig. 2) and an expression profile similar to those of some other *Tetrahymena* Nups (Fig. 5A). Localization of a GFP-fusion to NPCs was highly biased, albeit not exclusive, to the MAC (Fig. 5C). We therefore named this protein *Tt*Nup88, which is known to localize to the cytoplasmic side of the NPC in other species (Fig. 5B). As Nup88 in other species is known to interact with Nup214 and Nup98 (Fornerod et al., 1997), *Tt*Nup88 may contribute to the nucleus-specific localization of Nup214 and Nup98 paralogs.

**Fig. 5.**
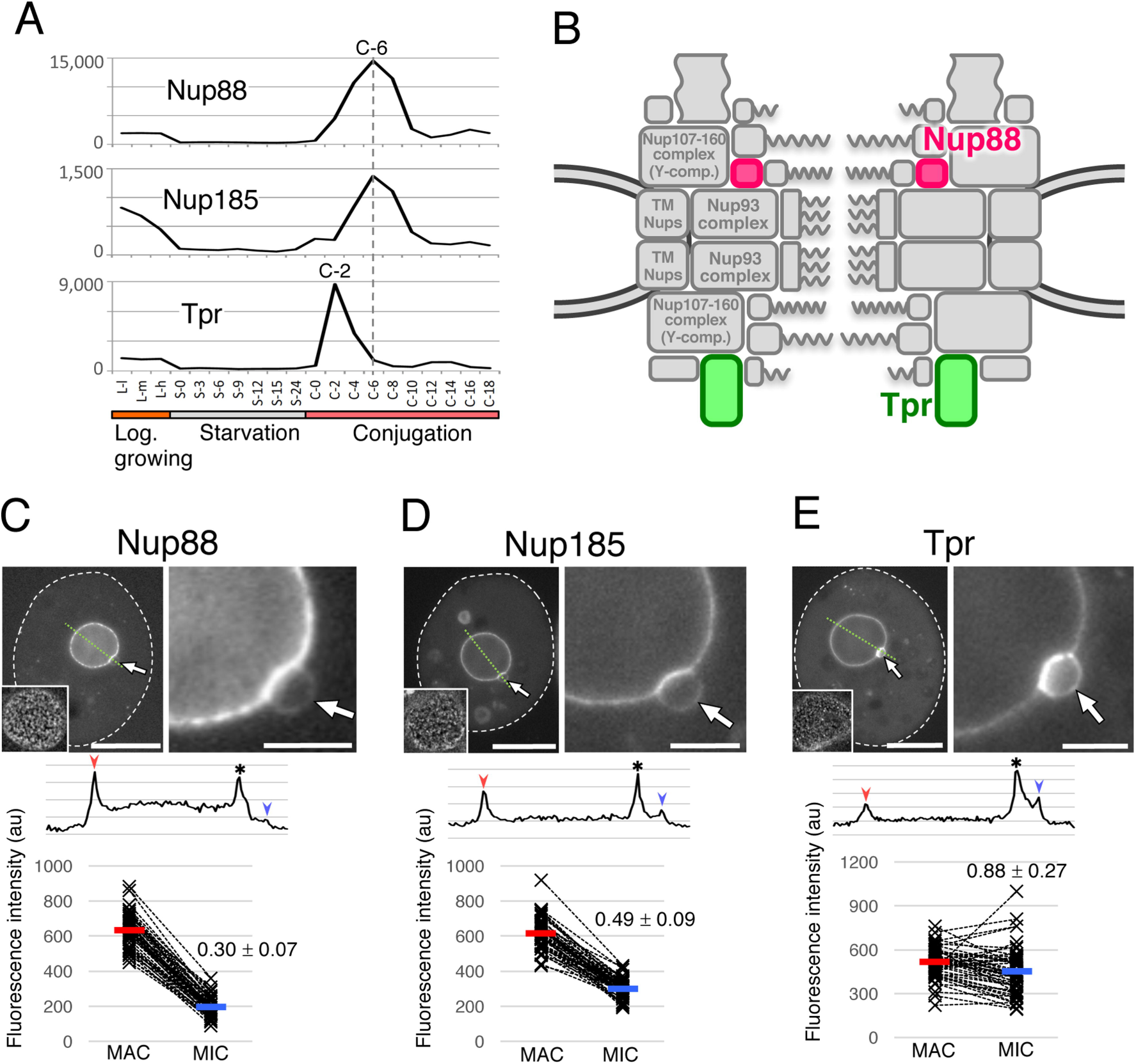
Nuclear localization and expression profiles of Nup88, Nup185, and Tpr. (A) Expression profiles. (B) The predicted positions of *Tt*Nup88 and *Tt*Tpr. The position of Nup185 is unknown. (C) The subcellular localization of ectopically expressed GFP**–***Tt*Nup88. The left panel shows a whole cell, and its nuclear region is enlarged in the right panel. White broken lines represent the borders of cells. Inset in the left panel shows the deconvoluted image focused on the MAC surface. Arrows indicate the position of the MICs. Bars indicate 20 μm for the left panel and 5 μm for the right panel. A line profile and plots of fluorescence intensity are shown under each image panel, as in Fig. 3B. The fluorescence intensity of the MIC NE is significantly lower than that of the MAC NE (*P* < 10^−39^). (D) Subcellular localization of ectopically expressed GFP**–**Nup185. The fluorescence intensity of the MIC NE is significantly lower than that of the MAC NE (*P* < 10^−30^). (E) Subcellular localization of ectopically expressed GFP**–***Tt*Tpr. The fluorescence intensity of the MIC NE is slightly lower than that of the MAC NE (*P* = 0.0024, by Student’s *t*-test).

TTHERM_00755920 (encoding a 185 kDa protein), which lies adjacent to the open reading frame (ORF) of MacNup214, attracted our interest because its predicted molecular structure resembled those of large scaffold Nups such as Nup160, Nup155, and Nup133, and because its expression profile is similar to those of some other *Tetrahymena* Nups (Fig. 5A). A GFP-fusion localized to NPCs, with a bias to the MAC (Fig. 5D). Based on its predicted molecular weight, we named this protein Nup185. Nup185 contains a conserved domain ‘Nucleoporin (SSF117289)’ (Fig. 2), which is generally found near the N-terminal regions of Nup155 and Nup133 homologs. The expression peak of Nup185 appeared at C-6 (Fig. 5A).

To assess the location of Nup185 within the NPC architecture, we identified interacting proteins by immunoprecipitating GFP–Nup185. One interacting protein was TTHERM_00268040, which bears predicted coiled-coil motifs throughout its entire sequence (Fig. 2) and is thus similar to the nuclear basket component, Tpr (Fig. 5B). TTHERM_00268040 fused with GFP localized equivalently to MAC and MIC NPCs (Fig. 5E). This protein is a likely ortholog of human Tpr; therefore, we named it *Tt*Tpr. Nup185 did not interact with any members of the Y- or Nup93 complexes (Table S6).

### The transmembrane Nups Pom121 and Pom82 show nucleus-specific localization

Some but not all of the transmembrane (TM) Nups are conserved between vertebrates and yeasts: the former have POM121, gp210, and NDC1 (Cronshaw et al, 2002; Stavru et al, 2006), while the latter have Pom34, Pom152, and Ndc1 (Rout et al, 2000; Asakawa et al, 2014). The only reported TM Nup in *T. thermophila* is gp210 (Iwamoto et al., 2009). Because all *Tetrahymena* Nups identified so far have a similar expression pattern, in which a large expression peak appears during early conjugation stage (Figs 3C, 4C and 5A), we used expression profiling and TM domain search to identify possible TM Nups in the updated TetraFGD and the TMHMM Server (http://www.cbs.dtu.dk/services/TMHMM-2.0/), respectively. Using this approach, we found two candidate TM Nups. Each has one TM domain and an FG-repeat region (“*Tt*Pom121” and “*Tt*Pom82” in Fig. 6A). Their expression profiles are shown in Fig. 6B.

One of the TM Nup candidates (TTHERM_00312730; *Tt*Pom121) has an N-terminal TM domain and C-terminal FG repeats (Fig. 6A, middle) with a deduced molecular weight of 129 kDa. These attributes are very similar to those of vertebrate POM121 (compare top and middle parts of Fig. 6A) (Rothballer and Kutay, 2012). *Tt*Pom121 fused with GFP at its C-terminus (*Tt*Pom121–GFP) localized specifically to MAC NPCs (Fig. 6C, upper). Consequently, this protein is the likely *Tetrahymena* ortholog to human POM121; therefore, we named it *Tt*Pom121.

Notably, when GFP was fused with the N-terminus of *Tt*Pom121 at a region close to the TM domain (GFP-*Tt*Pom121), the tagged protein localized in the MAC nucleoplasm, but not in MAC NPCs or the MIC nucleoplasm (Fig. 6C, lower). This result suggests that *Tt*Pom121 bears a MAC-specific nuclear localization signal (NLS) in its N-terminal region. Similarly, POM121 homologs in vertebrates have NLS sequences in the N-terminal region (Yavuz et al., 2010; Funakoshi et al., 2011).

In contrast, the other TM Nup candidate (TTHERM_00375160; TtPom82) localized exclusively to MIC NPCs (Fig. 6D, upper). This protein has predicted molecular features that have not been reported in Nups from any other organism: a TM domain near the C-terminus, central coiled-coil, and N-terminal FG repeats (Fig. 6A, bottom). We named this protein *Tt*Pom82 according to its predicted molecular weight (82 kDa). A construct lacking the TM domain showed diffuse cytoplasmic localization (Fig. 6D, lower), suggesting that MIC NPC-specific localization of *Tt*Pom82 does not depend on the MIC-specific nuclear transport of *Tt*Pom82. This result suggests that *Tt*Pom121 and *Tt*Pom82 use different mechanisms to target to the MAC and MIC NPCs.

Next, we performed immuno-electron microscopy (iEM) for the Pom proteins using anti-GFP antibody in order to know their sub-NPC localization. Intriguingly, their sub-NPC localizations were opposite; Pom121 was exclusively localized to the nuclear side of the MAC NPC (Fig. 6E), whereas Pom82 was exclusively localized to the cytoplasmic side of the MIC NPC (Fig. 6F).

**Fig. 6.**
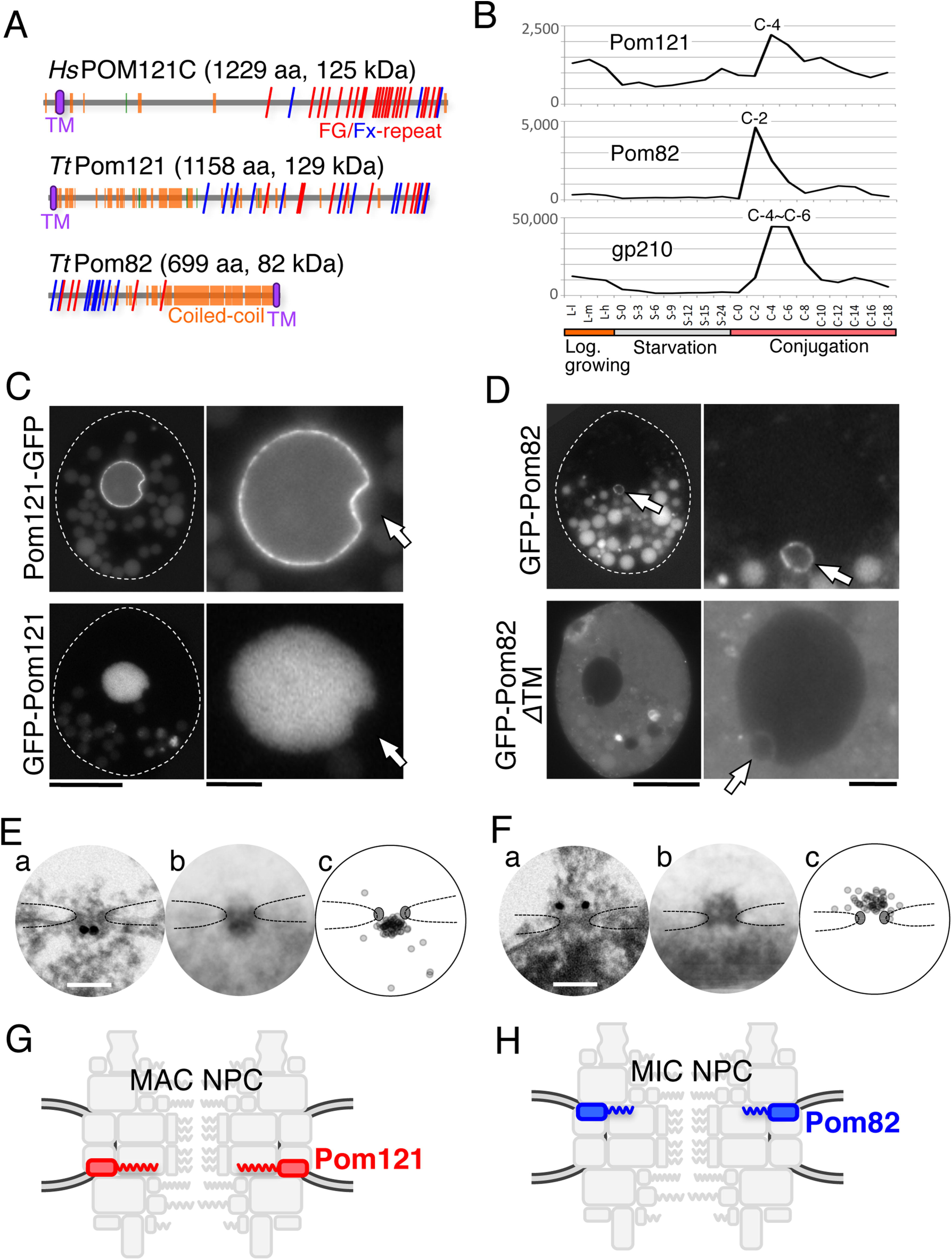
Two novel pore membrane proteins show nuclear specificity. (A) Illustration of molecular profiles. The frequency and positions of FG repeats are compared between *T. thermophila* Pom proteins and human POM121C (accession: A8CG34). Red and blue slanting lines represent FG and FX (X means any amino acid residue, but the majority are N, Q, and S), respectively. Orange and green boxes represent α-helices and β-strands, respectively. Purple ellipses represent predicted TM domains. (B) The expression profiles of nucleus-specific Pom-s and shared *Tt*gp210, as in Fig. 3C. (C) Fluorescence micrographs show ectopically expressed GFP-tagged *Tt*Pom121. Left panels show whole cells, and the right panels show enlarged images of the nuclear regions. White broken lines represent the borders of cells. Arrows indicate the position of MICs. Bars indicate 20 μm for the left panels and 5 μm for the right panels. (D) Fluorescence micrographs show GFP-tagged Pom82 (full length, 1**–**699 aa) and GFP**–**Pom82ΔTM (transmembrane domain-deletion mutant, 1**–**678 aa) both ectopically expressed. Arrows indicate the position of the MICs. Other fluorescent bodies dispersed in the cytoplasm are phagosomes taking in materials derived from the culture medium. (E, F) iEM for Pom121-GFP localizing to the MAC NPC (E) and GFP-Pom82 localizing to the MIC NPC (F) using anti-GFP antibody. (a) Immuno-electron micrographs for a single NPC. Dark dots represent signals of gold particles. Bars, 100 nm. (b) Images present a projection image of 20 immuno-electron micrographs of NPCs decorated with gold particles. (c) The positions of individual gold particles in (b) are plotted. Broken lines trace nuclear envelope, and upper and lower sides are cytoplasm and nucleoplasm, respectively. (G) The position of *Tt*Pom121 within the MAC NPC architecture. (H) The position of *Tt*Pom82 within the MIC NPC architecture.

Given the difference in molecular features, their behaviors when the TM domain function was disrupted, and their sub-NPC localizations, Pom121 and Pom82 are unlikely to be functional homologs of each other. Taken together, these findings lead to the conclusion that MAC and MIC NPCs contain distinct TM components (Fig. 6G,H). The protein components of MAC and MIC NPCs are summarized in Fig. 7.

**Fig. 7.**
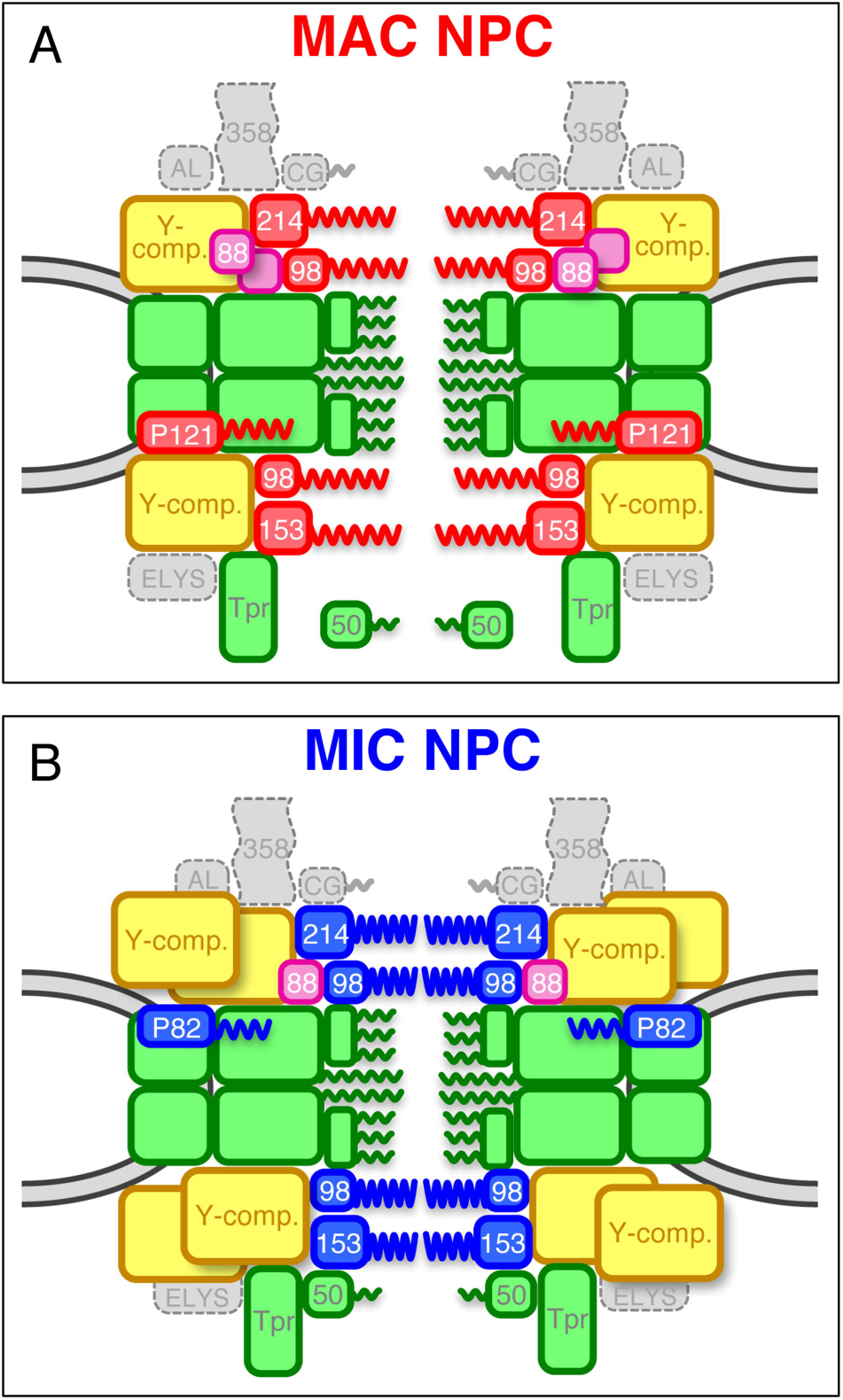
Schematic models of MAC and MIC NPCs. (A) Deduced composition of the MAC NPC. (B) Deduced composition of the MIC NPC. Boxes colored in red and blue represent MAC-specific and MIC-specific components, respectively: Pom121 (P121), Pom82 (P82), Nup98 paralogs (98), Nup214 (214), and Nup153 (153). Green boxes represent shared components including the nuclear basket structure Tpr and its associated Nup50 (50). *Tt*Nup50 is distributed mostly in the nucleoplasm in MACs, whereas it localizes to the NPC in MICs (Malone et al., 2008; Iwamoto et al., 2009). Yellow boxes are MIC-biased Y-complexes, and purple boxes are MAC-biased *Tt*Nup88 (88). The number of duplications of yellow and purple boxes does not reflect the actual quantity of those components *in vivo.* Homologs of Nup358 (358), hCG1 (CG), ALADIN (AL), and ELYS constituting the cytoplasmic structure, were not found in *T. thermophila*.

One TM Nup, found in both fungi and animals but missing from our *Tetrahymena* catalog, is Ndc1. We identified a potential Ndc1 homolog in TTHERM_00572170, a protein with six predicted TM domains that is co-transcribed with other Nups (see http://tfgd.ihb.ac.cn/search/detail/gene/TTHERM_00572170). However, neither N- nor C-terminal GFP fusions of this protein localized to NPCs (Fig. S3D). Therefore, *Tetrahymena* NPCs may lack Ndc1. Similarly, Ndc1 has not been detected in *Trypanosoma* NPCs (Obado et al., 2016).

### The permeability of the nuclear pore differs between MAC and MIC

To better understand the functional consequences of structural differences between MAC and MIC NPCs, we examined the relative pore exclusion sizes by asking whether probes of different sizes could gain access to each nucleoplasm. GFP (approx. 28 kDa) was excluded only from MICs, whereas GFP–GST (more than 100 kDa due to its oligomerization) was excluded from both MACs and MICs (Fig. S4A). In addition, FITC-dextran of 40 kDa could enter MACs, whereas 70-kDa FITC-dextran was completely excluded (Fig. S4B). These results indicate that MAC pores exclude molecules greater than approximately 50 kDa, which is similar to the permeability size limit of nuclear pores in other species (Paine et al., 1975; Gorlich and Mattaj, 1996; Keminer and Peters, 1999). On the other hand, MIC pores impose a much smaller exclusion size, and exclude molecules of even 10–20 kDa (Fig. S4B). This difference in exclusion size may be due to differences between the protein composition and structural arrangement of NPCs of these dimorphic nuclei.

## DISCUSSION

We have now identified 28 nucleoporins in the ciliate *T. thermophila*: 15 Nups reported here, and 13 in our previous study (Iwamoto et al., 2009). This total comprises 24 different Nups for the MAC and MIC: this number includes 18 Nups commonly localized in both nuclei, 4 Nups with nucleus-specific homologues (Nup214, Nup153, Nup98A, and Nup98B), and *Tt*Pom82 and *Tt*Pom121. This total is somewhat smaller than the roughly 30 Nups known in other eukaryotes, e.g. 34 in human and in *Drosophila melanogaster*, 27 in *Caenorhabditis elegans,* 33 in *S. pombe,* and 35 in *S. cerevisiae* (Rothballer and Kutay, 2012; Asakawa et al., 2014). The deficit in *T. thermophila* Nups is due to the absence of homologs for Nup358, GLE1, hCG1/Nup42, Nup43, Nup37, Centrin-2, Nup53, TMEM33, ELYS, and Aladin. Similarly, the protist *Trypanosoma brucei* is missing Nup358, GLE1, hCG1/Nup42, Nup37, Centrin-2, TMEM33, and ELYS, and 25 Nups in total have been identified by interactome analysis (DeGrasse et al, 2009; Obado et al., 2016). One conserved Nup identified in *Trypanosoma* but not *Tetrahymena* is Nup53 (*Tb*Nup65 (XP_822630.1)) (Obado et al., 2016). This raises the question of whether a *T. thermophila* Nup53 homolog eluded our search due to sequence or structural divergence. Alternatively, *T. thermophila* may have lost a Nup that is not essential for viability.

### A role of nucleus-specific Nups

We previously reported that the GLFG-repeat and NIFN-repeat domains in MacNup98s and MicNup98s, respectively, are involved in the nucleus-specific transport of linker histones (histone H1 and MLH, respectively), arguing that these nucleus-specific Nups are determinants of nucleus-specific transport (Iwamoto et al., 2009). Importantly, we can now expand this argument, since our expanded catalog shows that all NPC subunits that are nucleus-specific are FG-Nups: Nup214, Nup153, Nup98 and Pom-s. Since the FG-repeats interact with nuclear transport receptors such as importin-β family proteins (Allen et al., 2001; Isgro and Schulten, 2005; Liu and Stewart, 2005; Tetenbaum-Novatt et al., 2012), specificity for the MAC or MIC is likely to be determined in cooperation with importin-βs. This idea is also supported by the presence of nucleus-specific importin family proteins (Malone et al., 2008).

It is interesting to note that both MAC- and MIC-specific Nups contain atypical repeat motifs: NIFN, but also more subtle variations on the FG-repeat: FN, FQ, FA, FS and so on (Fig. 2). Because the NIFN-repeat domain of MicNup98A is known to function in blocking misdirected nuclear transport of MAC-specific linker histones (Iwamoto et al., 2009), the atypical FG-repeats may similarly be involved in controlling nucleus-specific transport of particular proteins. However, importin-βs that preferentially interact with the NIFN-repeat and their cargos have not been found, and thus the complete role of the NIFN-repeat motif in nucleus-specific transport remains to be elucidated.

### A role of biased Nups to build different NPC structures

The nucleus-specific Nups generate obvious structural differences between MAC and MIC NPCs. However, these different components have to be integrated into two NPC scaffold structures that are constructed of the same components. One way to make different structures from the same components may be to incorporate different amounts of these components, leading to different structures that allow biased localization/assembly of nucleus-specific components. The localization of the Y-complex (Fig. 3B) and Nup88 (Fig. 5C) was highly biased to either MICs or MACs, respectively. Thus, these biased components may be critical for directing assembly of MAC- or MIC-type NPCs. Consistent with this idea, Nup98 homologs in vertebrates interact with the Y-complex components Nup96 (Hodel et al., 2002) and Nup88 (Griffis et al., 2003). This model raises the question of how structurally similar paralogs in *Tetrahymena* can differentially recruit nucleus-specific FG-Nups.

The copy number of the Y-complex within individual NPCs differs between the MAC and MIC (Fig. 3B,D), indicating that at least two NPC structures with different Y-complex stoichiometries can form in ciliates. This quantitative difference in Y-complex incorporation may be directed by membrane Nups. The nucleus-specific TM Nups, Pom121 and Pom82, are currently strong candidates for initiating NPC assembly on the nuclear membrane. In vertebrates, Pom121 binds the Y-complex through a Nup160 homolog (Mitchell et al., 2010). In *Tetrahymena, Tt*Pom121 and *Tt*Pom82 may differentially affect Y-complex integration into MAC or MIC NPCs. This model can be extended to biased integration of Nup98 paralogs, since Pom121 has been shown to directly bind Nup98 (Mitchell et al., 2010), supporting our idea that biased Nups and nucleus-specific Nup98 paralogs cooperate to build two distinct NPCs. In this model, the acquisition of specialized Pom proteins might have been one of the most crucial evolutionary events for generating nuclear dimorphism in ciliates. Taken overall, our study contributes to understanding the diversity of NPC architectures in eukaryotes, including potential functional and evolutionary aspects.

## MATERIALS AND METHODS

### *In silico* genomic database analysis and secondary structure prediction

We searched for candidates Nups using protein BLAST on the NCBI website and *Tetrahymena* Genome Database Wiki (http://ciliate.org/index.php/home/welcome) (Eisen et al., 2006; Stover et al., 2012). Expression profiles based on microarray data (http://tfgd.ihb.ac.cn/tool/exp) were obtained from the TetraFGD (http://tfgd.ihb.ac.cn/) (Miao et al., 2009). We identified the candidate proteins as Nups when the expression profile satisfied two conditions: First, the amount of expression is lower in vegetative stages than in conjugation stages. Second, expression peaks appear in between C-2 and C-8 stages of conjugation. Secondary structures and transmembrane domains were predicted by PSIPRED (http://bioinf.cs.ucl.ac.uk/psipred/) and the TMHMM Server (http://www.cbs.dtu.dk/services/TMHMM-2.0/), respectively. Coiled-coil regions were predicted by PBIL Coiled-Coils prediction (https://npsa-prabi.ibcp.fr/cgi-bin/npsa_automat.pl?page=npsa_lupas.html) or SIB COILS (http://embnet.vital-it.ch/software/COILS_form.html). Conserved domains were searched for using InterPro (http://www.ebi.ac.uk/interpro/).

### DNA construction

cDNAs were amplified by PrimeSTAR reagent (Takara, Kyoto, Japan) from the reverse transcripts prepared from the total RNA fraction of vegetative or conjugating cells as described previously (Iwamoto et al., 2009). The cDNAs were digested with *Xho*I and *Apa*I, and cloned into the pIGF1 vector to ectopically express them as N-terminal GFP-tagged proteins (Malone et al., 2005). The pIGF1C vector with the multi-cloning site at the 5’ site of the GFP-coding sequence was generated by modifying the pIGF1 vector, and used to ectopically express GFP-tagged Nup58 and Pom121 as C-terminal GFP-tagged proteins: the cDNAs of these Nups were cloned into the pIGFIC vector using the *Xho*I and *Kpn*I sites. To endogenously express Nups tagged with a fluorescent protein at the C-termini of the macronuclear ORFs, MicNup214, Nup160, and Nup133 were tagged with GFP using a pEGFP-neo4 vector (Mochizuki, 2008) (a kind gift from Dr. K. Mochizuki, IMBA), MicNup153 was tagged with mNeon using a p2xmNeon_6xmyc_Neo4 vector (a kind gift from Dr. Turkewitz, Univ. of Chicago), and Sehl was tagged with mCherry using a pmCherry-pur4 vector (Iwamoto et al., 2014). Primers used in this study are listed in Table S7.

### Expression of GFP-Nups in *Tetrahymena* cells

Conjugating cells were subjected to transfection by electroporation using a Gene Pulser II (Bio-Rad, Hercules, CA) as described previously (Iwamoto et al., 2014; 2015). The resulting cell suspension was cultivated for 18 h and then treated with selection drugs, paromomycin sulfate (Sigma-Aldrich, St. Louis, MO) at 120 μg/ml when using pIGF1, pIGF1C, pEGFP-neo4, and p2xmNeon_6xmyc_Neo4 vectors, or puromycin dihydrochloride (Fermentek, Jerusalem, Israel) at 200 μg/ml when using a pmCherry-pur4 vector. Cadmium chloride was also added at 0.5 μg/ml to induce the expression of drug resistant genes for pEGFP-neo4, p2xmNeon_6xmyc_Neo4, and pmCherry-pur4 vectors. Resistant cells usually appeared within a few days after the drug was added. We checked that at least 5 independent clones (*i.e*., grown in 5 different wells) exhibited the same intracellular localization of each GFP–Nup.

### Immunoprecipitation

For immunoprecipitation, GFP–Nup-expressing cells in logarithmic growth were pretreated with 0.5 mM PMSF for 30 min at 30°C and then collected by centrifugation. The cells were resuspended at 2.5 × 10^6^ cells/ml in homogenization buffer composed of 150 mM NaCl, 1% Triton X-100, 2 mM PMSF, and Complete Protease Inhibitor Cocktail (Roche Diagnostics, Mannheim, Germany), and then homogenized with sonication on ice. The supernatant obtained after centrifugation at 10,000×g for 15 min was pretreated with Protein A Sepharose to absorb non-specifically bound proteins. After removal of the beads by low-speed centrifugation, the supernatant was incubated with 50 μg anti-GFP rabbit polyclonal antibody (#600-401-215, Rockland Immunochemicals, Limerick, PA) for 2 h at 4°C. To collect immunoprecipitated target proteins of interest, fresh Protein A Sepharose was added, incubated for another 2 h at 4°C, and then collected by centrifugation. After brief washing with homogenization buffer, the Sepharose beads were incubated with NuPAGE sample buffer (Thermo Fisher Scientific, Waltham, MA) to elute bound proteins. The proteins were separated by SDS-PAGE.

### Mass-spectrometry analysis

The gel sample lane was cut into several pieces, and each treated with trypsin. The trypsinized peptide sample was subjected to liquid chromatography/tandem mass spectrometry (LC/MS/MS) using the LXQ linear ion trap (Thermo Finnigan, San Jose, CA) equipped with a Magic2002 and nanospray electrospray ionization device (Michrom BioResources, Auburn, CA and AMR, Tokyo, Japan), as described previously (Obuse et al., 2004). The LC-MS/MS data were searched by Mascot (Matrix Science, London, UK) with a non-redundant *T. thermophila* specific database (25,131 sequences) constructed from the nr NCBI database. The resulting files were loaded into Scaffold software (Proteome Software, Portland, OR) for comparing identified proteins between samples.

### Microscopic observation

Intracellular localizations of GFP-tagged Nups were observed by fluorescence microscopy (IX-70; Olympus, Tokyo, Japan). Images were taken using the DeltaVision microscope system (GE Healthcare, Issaquah, WA) with oil-immersion objective lens UApo40 (NA=1.35) (Olympus). Line profiles of fluorescence intensity were obtained using a measurement tool included in the DeltaVision system. Background fluorescence measured cytoplasm as an averaged value of 5×5 pixels was subtracted from the peak values of fluorescence on the NE.

### Indirect Immunofluorescence staining

*Tetrahymena* cells expressing GFP-tagged Nups were first fixed with cold methanol for 20 min, and then additionally fixed with 4% formaldehyde in PBS for 20 min. After treated with 1% bovine serum albumin (BSA), cells were treated with 5 μg/ml anti-GLFG monoclonal antibody 21A10 for 2-3 hrs (Iwamoto et al., 2013). After washing with PBS, cells were treated with Alexa Fluor 594-conjugated goat anti-mouse IgG at 1/1000 dilution for 1 h (Thermo Fisher Scientific). Images of forty z-sections with a 0.2-μm interval were taken for cells using the DeltaVision microscope system with oil-immersion objective lens PlanApoN60OSC (NA=1.4) (Olympus), and were processed by deconvolution using SoftWoRx software equipped with the microscope.

### Immuno-electron microscopy

*Tetrahymena* cells expressing GFP-tagged Nups were fixed with 4% formaldehyde for 30 min. After washing 3 times with PBS, they were permeabilized with 0.1% saponin for 15 min at room temperature. After treatment with 1% BSA, cells were incubated with anti-GFP polyclonal antibody (Rockland Immunochemicals) at 1/200 dilution for 2 hrs, washed three times with PBS, then incubated with FluoroNanogold-anti rabbit Fab’ Alexa Fluor 594 (Nanoprobes, Yaphank, NY) at 1/400 dilution for 1 h. The immunolabelled cells were fixed with 2.5% (w/v) glutaraldehyde (Nacalai tesque, Kyoto, Japan) for 1 h. After washing with 50 mM HEPES (pH 5.8) they were incubated with silver enhancement reagent (Tange et al., 2016) for 7 min. The reaction was stopped by washing three times with distilled water. Then the cells were post-fixed with 1% OsO_4_ for 15 min, electron stained with 2% uranyl acetate for 1 h, dehydrated with sequentially increased concentrations of ethanol, and embedded in epoxy resin (Epon812). The ultrathin sections sliced from the resin block were stained with 4% uranyl acetate for 15 min and lead citrate (Sigma-Aldrich) for 1 min, and observed by a transmission electron microscope JEM-1400 (JEOL, Tokyo, Japan) with an acceleration voltage of 80 kV.

## Acknowledgements

We thank *Tetrahymena* Stock Center at Cornell University, *Tetrahymena* Functional Genomics Database, *Tetrahymena* Genome Database Wiki, Drs. K. Mochizuki and A.P. Turkewitz for providing materials or valuable information. We thank S. Shibata and N. Shirai for technical assistance of LC-MS/MS analysis. We also thank Drs. D.B. Alexander, H. Asakawa, A.P. Turkewitz, S.O. Obado and M.P. Rout for critical reading of this paper.

## Competing interests

No competing interests declared.

## Author contributions

MI, HO, CM, YF, and KN performed the experiments. MI, KN, CO, YH and TH designed the experiments. All authors examined and discussed the data, and MI, CO, YH, and TH wrote the manuscript.

## Funding

This work was supported by grants from the Japan Science and Technology Agency to TH and Japan Society for the Promotion of Science Kakenhi Grant Numbers JP24570227, JP15K07066 to MI, JP15H01462 to KN, JP20114006, JP25116004 to CO, JP26116511, JP16H01309, JP26251037 to YH, and JP23114724, JP26291007, JP25116006 to TH.

